# ^13^CO_2_ pulse labelling reveals species-specific alterations in carbon allocation and volatile organic compound emissions under heat stress

**DOI:** 10.64898/2026.08.24.746709

**Authors:** Stefanie Dumberger, Clara Stock, Mirjam Meischner, Melissa Wannenmacher, Helen Vogt, Phyllis Lua-Mellmann, Kathrin Kühnhammer, Jürgen Kreuzwieser, Christiane Werner, Simon Haberstroh

## Abstract

- Temperate forests increasingly face extreme air temperature, but plant physiological responses, particularly alterations in carbon allocation or protection via volatile organic compound (VOC) emissions, remain poorly understood.
- We pulse-labelled well-watered saplings of *Fagus sylvatica* and *Pseudotsuga menziesii* in a controlled heat stress experiment with ^13^CO_2_ to quantify heat-induced shifts in CO_2_, VOC and C pool exchange, specifically analyzing compound-specific δ^13^C of terpenoids, water-soluble organic matter (WSOM) and dark respiration.
- Under heat stress, up to 50% of fresh assimilates were directed to maintenance respiration and 1-2% to VOC emissions, while net assimilation and water use efficiency decreased by 50-75% in both species. Heat directly affected metabolic processes and reduced turnover rates of fresh assimilates in *F. sylvatica*, but accelerated them in *P. menziesii.* Strong ^13^C labelling of some compounds, particularly acyclic ones, suggested increased *de novo* synthesis of specific terpenoids for heat stress protection.
- By tracing the fate of recently assimilated ^13^CO_2_ we demonstrate that heat stress reduces net carbon uptake and water use efficiency, disrupts turnover of C pools and increases carbon loss via respiration and *de novo* synthesis of specific VOCs, potentially diminishing net carbon uptake of forests under future heat extremes.

## Introduction

Globally, forests are suffering from increased compound droughts, i.e., simultaneously occurring edaphic and atmospheric drought, caused by a lack of precipitation and high air temperature (Bastos et al., 2023; Dunn et al., 2024; Fang et al., 2022; Werner et al., 2025). Especially ecosystems of the temperate zone, such as in Central Europe, are prone to the occurrence of compound droughts with devastating effects on ecosystem functioning and survival (Haberstroh et al., 2022; Knutzen et al., 2025; Schuldt et al., 2020). Yet, it remains challenging to disentangle the impacts of edaphic and atmospheric drought, since natural heat waves usually co-occur with elevated vapor pressure deficit and reduced precipitation (De Boeck et al., 2010). Moreover, physiological responses, such as stomatal closure, reduced growth, leaf shedding or tree mortality, are often comparable (Haberstroh et al., 2025; Marchin et al., 2022; Teskey et al., 2015; Urban et al., 2017). Studies on edaphic drought have dominated the research landscape in recent years, while the isolated impact of atmospheric drought caused by high air temperature remains less well understood (Grossiord et al., 2020; He et al., 2022; Mirabel et al., 2023; Novick et al., 2024; Yuan et al., 2025).

Plants operate at thermal optima and physiological activity is impaired, if air temperature is excessively high (Berry & Bjorkman, 1980; Niinemets, 2018; Teskey et al., 2015) or low (Berry & Bjorkman, 1980; Sage & Kubien, 2007). Beyond the thermal optimum, carbon assimilation is compromised, which can be attributed to the inactivation of Rubisco by decreased enzymatic activity of Rubisco activase, increased photorespiration, inhibition of photosystem II and the disruption of thylakoid membranes by formation of reactive oxygen species (ROS) and increased membrane fluidity (Hüve et al., 2011; Niinemets, 2018; Teskey et al., 2015). While the constrained carbon uptake under heat exposure is well described, our knowledge about plant-internal carbon allocation patterns in response to stress conditions is scarce (Cabon, 2026; Gessler & Zweifel, 2024; Trugman & Anderegg, 2025; Werner et al., 2020).

Plants have evolved a plethora of protective mechanisms to buffer the negative effects of heat exposure, including stomatal aperture for evaporative cooling (Diao et al., 2024; Drake et al., 2018; Larcher, 2003; Urban et al., 2017), increased dark respiration (Niu et al., 2024; Scafaro et al., 2021), emission of volatile organic compounds (VOCs), (Delfine et al., 2000; Loreto et al., 1998; Sharkey et al., 2001), adaptation of leaf osmotic potential and membrane fatty acid saturation (Hüve et al., 2012; Tarvainen et al., 2022), and the production of heat shock proteins (Jagadish et al., 2021; Vierling, 1991). Under well-watered conditions, evaporative cooling of leaves can critically enhance the thermal safety margin, but also increases water loss (Bachofen et al., 2025; Meischner et al., 2024) resulting in a decreased water use efficiency and increased depletion of water resources (Diao et al., 2024; Urban et al., 2017). Stimulation of respiration is related to stabilization, repair and novel production of enzymes and cell membranes, increased utilization of starch due to reduced sucrose synthesis and the activation of ROS scavenging pathways (Scafaro et al., 2021; Teskey et al., 2015). Increased day- and nighttime respiration constraints already reduced carbon uptake resulting in a negative carbon balance under severe heat stress (Rehschuh et al., 2022; Werner et al., 2020). Emission of VOCs, particularly terpenoids and isoprene, is considered as a key protective mechanism against exposure to heat and excessive radiation (Loreto & Schnitzler, 2010; Peñuelas & Llusià, 2003; Vickers et al., 2009). While high air temperature increases volatility (Guenther, 2013; Holopainen et al., 2018; Makkonen et al., 2012), it can also stimulate production and emission rates of VOCs (Bourtsoukidis et al., 2024; Haberstroh et al., 2018; Meischner et al., 2024; Staudt & Bertin, 1998). Yet, temperature-dependent emission differs between species with specialized storage organs for VOCs (such as conifers) emitting a mixture of stored and *de novo* compounds and species without such features (e.g. a variety of broad-leaved species) depending primarily on *de novo* synthesis and freshly assimilated carbon (Daber et al., 2025; Kleist et al., 2012; Llusia et al., 2013; Staudt et al., 2017; Yáñez-Serrano et al., 2019).

The study of carbon stable isotope ratios (^13^C/^12^C) in combination with isotopic labelling approaches may elucidate the fate and pathways of freshly assimilated carbon (Epron et al., 2012; Werner et al., 2012), including the incorporation into *de novo* synthesized VOCs (Daber et al., 2025; Ghirardo et al., 2010; Haberstroh et al., 2019; Harley et al., 2014; Kleist et al., 2012; Kreuzwieser et al., 2021; Meischner et al., 2026; Werner et al., 2021). Moreover, isotopic labelling may also inform about carbon source and sink dynamics under stress conditions, particularly about the role of non-structural carbohydrates (NSCs) as a transient storage and buffer compartment (Blessing et al., 2015; Cabon, 2026; Gessler & Zweifel, 2024; Huang et al., 2024; Keel et al., 2006). NSCs comprise of two pools with high interconvertibility: non-soluble starch forms a transitory storage with frequent depletion, while water-soluble organic matter (WSOM) remains above a minimum threshold and serves more immediate functions, such as transport, osmolytes, substrate, respiration and defense (Hartmann & Trumbore, 2016; Huang et al., 2018; Martínez-Vilalta et al., 2016). Yet, knowledge of NSC flux dynamics under environmental stress is scarce, but may crucially contribute to our understanding of the coordination between carbon supply and demand (Cabon, 2026; Gessler & Zweifel, 2024; Trugman & Anderegg, 2025).

In Central European forests, deciduous European beech (*Fagus sylvatica* L.) and coniferous Douglas fir (*Pseudotsuga menziesii* (MIRB.) FRANCO) are two economically important tree species with differing properties regarding water use strategies, i.e. more anisohydric vs. more isohydric (Paligi et al., 2025; Schumann et al., 2024) and VOC storage, i.e. non-storing vs. storing species (Dindorf et al., 2006; Holzke et al., 2006; Lerdau et al., 1995; Pressley et al., 2004). Earlier work classified *P. menziesii* as well adapted to climate change, particularly in terms of drought-resistance (Lévesque et al., 2014; Ruehr et al., 2016; Spellmann et al., 2015), however, more recent studies suggested a high susceptibility to high air temperature and edaphic drought with significant reductions in productivity (Duarte et al., 2016; Enderle et al., 2024; Kunert et al., 2022; Leuschner & Meinzer, 2024). Heat and drought tolerance of *F. sylvatica* is medium compared to other broadleaved species such as *Quercus* and *Acer* spp. (Hauck et al., 2025; Münchinger et al., 2023) and it is suspected to be susceptible to future, hotter climate conditions (Deluigi et al., 2025; Geßler et al., 2007; Leuschner, 2020). However, in response to global warming growth was more impaired in *P. menziesii* than in *F. sylvatica* (Enderle et al., 2024).

The main objective of this study is to investigate the impact of high air temperature on gas exchange and carbon allocation in *F. sylvatica* and *P. menziesii* saplings by applying a ^13^CO_2_ pulse label. We hypothesize that heat stress (i) strongly increases transpiration and maintenance respiration and simultaneously decreases net carbon assimilation resulting in reduced water use efficiency, (ii) increases mean residence time of fresh assimilates due to decreased metabolic activity, and (iii) alters VOC emission rates and terpenoid composition. Specifically, we expect that elevated terpenoid emissions are mainly driven by increased *de novo* synthesis in *F. sylvatica* and increased release from storage in *P. menziesii*, while composition is altered by *de novo* synthesis of specific terpenoids to enhance thermotolerance.

## Material and Methods

### Plant material

The experiment was performed on two to four year-old European beech (*Fagus sylvatica* L.) and Douglas fir (*Pseudotsuga menziiesi* (MIRB.) FRANCO) saplings from nurseries in Southern Germany (Baumschulen Haage GmbH & Co. KG, Leipheim, Germany and Gustav Burger Forstbaumschulen, Zell am Harmersbach, Germany). From April to June 2023, plants were potted in boxes with a volume of 38 L filled with a 2 cm layer of expanded clay and a 5:1 mixture of soil (20% peat, Breisgau Kompost, Müllheim, Germany) and sand (0.7 -1.25 mm) with additional fertilizer (2 g L^-1^ NPK, Osmocote Exact Standard, Everris GmbH, Nordhorn, Germany). In each box, two saplings were growing together in both, inter- and intraspecific, combinations. The boxes were placed at the Chair of Ecosystem Physiology, University of Freiburg, on a scaffold covered with transparent foil to exclude natural precipitation and allow for controlled water supply.

### Experimental design and ecophysiological measurements

In July 2024, a total of 24 saplings were transferred into walk-in climate chambers (ThermoTec, Weilburg, Germany) and acclimatized for four days to a daily rhythm of 15 hours of daylight (25°C) with a photosynthetic photon flux density (PPFD) of 600 μmol m^-2^ s^-1^, seven hours of night-time conditions (15°C) and one hour of transient dusk and dawn, respectively. Relative humidity (RH) of ambient air was kept constant at 60% and soil moisture at 23.0 ± 0.1 Vol.% which was monitored by GS1 and 5TM soil moisture sensors (METER group, Munich, Germany).

After acclimatization, one branch of each sapling was placed in self-built enclosures (3 to 5 L) made of PET foil (Toppits, Cofresco Frischhalteprodukte GmbH & Co. KG, Minden), sealed airtight with plastic sealing tape (TEROSON RB II, Henkel AG, Düsseldorf, Germany) and rubber band and connected to an automated flow-through gas exchange measurement system (Daber et al., 2025; Fasbender et al., 2018; Meischner et al., 2026; Werner et al., 2020). Gas mixing inside the enclosures was ensured by small fans (MC25101V2-000U-A99, Sunon, Kaohsiung City, Taiwan). A zero-air generator (ZA-FID-AIR, LNI Swissgas GmbH, Kamen, Germany) provided VOC-free air through PFA-tubing (1/4”, Wolf-Technik eK, Stuttgart, Germany) with a controlled CO_2_ concentration of 470.8 ± 0.7 ppm and a ẟ^13^C ratio of -21.0 ± 0.9 ‰ to the cuvettes. ẟ^13^C ratio was depleted compared to ambient air due to the use of anthropogenic CO_2_ from combustion processes. Inflow was constant at a rate of 700 ml min^-1^ regulated by mass flow controllers (FMA5400A/5500A Series, Omega Engineering Inc., Deckenpfronn, Germany & 2SLPM Flow Controller, Alicat Scientific, Tucson, AZ, USA). One enclosure in each climate chamber was left empty as a reference for incoming air. Measurement air was led through PFA-tubing to the analyzer unit consisting of a non-dispersive infrared gas analyzer (LI-850, LI-COR Environmental, Bad Homburg, Germany), an isotope ratio infrared spectrometer (Delta Ray IRIS, Thermo Fisher Scientific, Darmstadt, Germany) and a proton-transfer-reaction time-of-flight mass spectrometer (PTR-ToF-MS 4000 ultra, Ionicon Analytic, Innsbruck, Austria) measuring ^12^CO_2_, ^13^CO_2_, H_2_O and BVOC concentrations at a resolution of 10 sec, 5 sec and 20 sec, respectively. Switching between the enclosures was realized every 6 minutes by a multi-position valve (Valco Instruments Inc., Schenken, Switzerland). Data acquired during the first day of the continuous measurements was discarded to avoid interferences caused by the installation of the enclosures (Duhl et al., 2008; Meischner et al., 2024). Diurnal net assimilation of CO_2_ (*A_net_*), nocturnal respiration (*R_n_*), transpiration (*E*), stomatal conductance (*g_s_*) and water use efficiency (*WUE*) were calculated from the gas concentration differences between the empty reference enclosures and the leaf enclosures (Supporting Information Eqn. S1-S4) according to von Caemmerer & Farquhar (1981). To determine leaf and projected needle area inside the cuvettes, leaves and needles were placed evenly on a commercial scanner after full plant harvest and scans were analyzed using the GSA Image Analyzer Software (GSA GmbH, Rostock, Germany).

After three days of pre-stress measurements, air temperature (*T_air_*) of one climate chamber was increased to 35 °C by a temperature ramp (2 °C h^-1^) and nocturnal *T_air_* was set to 25°C. After four days of heat stress, a ^13^CO_2_ pulse label was applied to all saplings (Figure 1). For four hours, enclosures were expanded to a volume of 6 to 8 L and approximately 50% of the entire leaf area was placed inside them. Supplied air was then enriched with 3.5 ml min^-1^ of pure ^13^CO_2_ increasing the isotopic ratio (ẟ^13^C) on average to +283.2 ± 25.7 ‰ (max. +751.3 ± 144.0 ‰) and the total CO_2_ concentration to 486.3 ± 1.9 ppm. After nine days of heat stress, *T_air_* was decreased from 35°C to 25°C by a temperature ramp (2°C h^-1^), enclosures were removed and saplings remained in the climate chambers for five more days to ensure full translocation of the ^13^CO_2_ label to its respective sink tissue. Thereafter, the entire biomass was harvested and separated into the foliage inside and outside the cuvettes, xylem and phloem tissue of stems and twigs and roots. For the sampling of phloem and xylem tissue, outer dead bark tissue was removed gently with a scapular, then the scapular was used to separate living phloem from woody xylem tissue.

**Figure 1:**
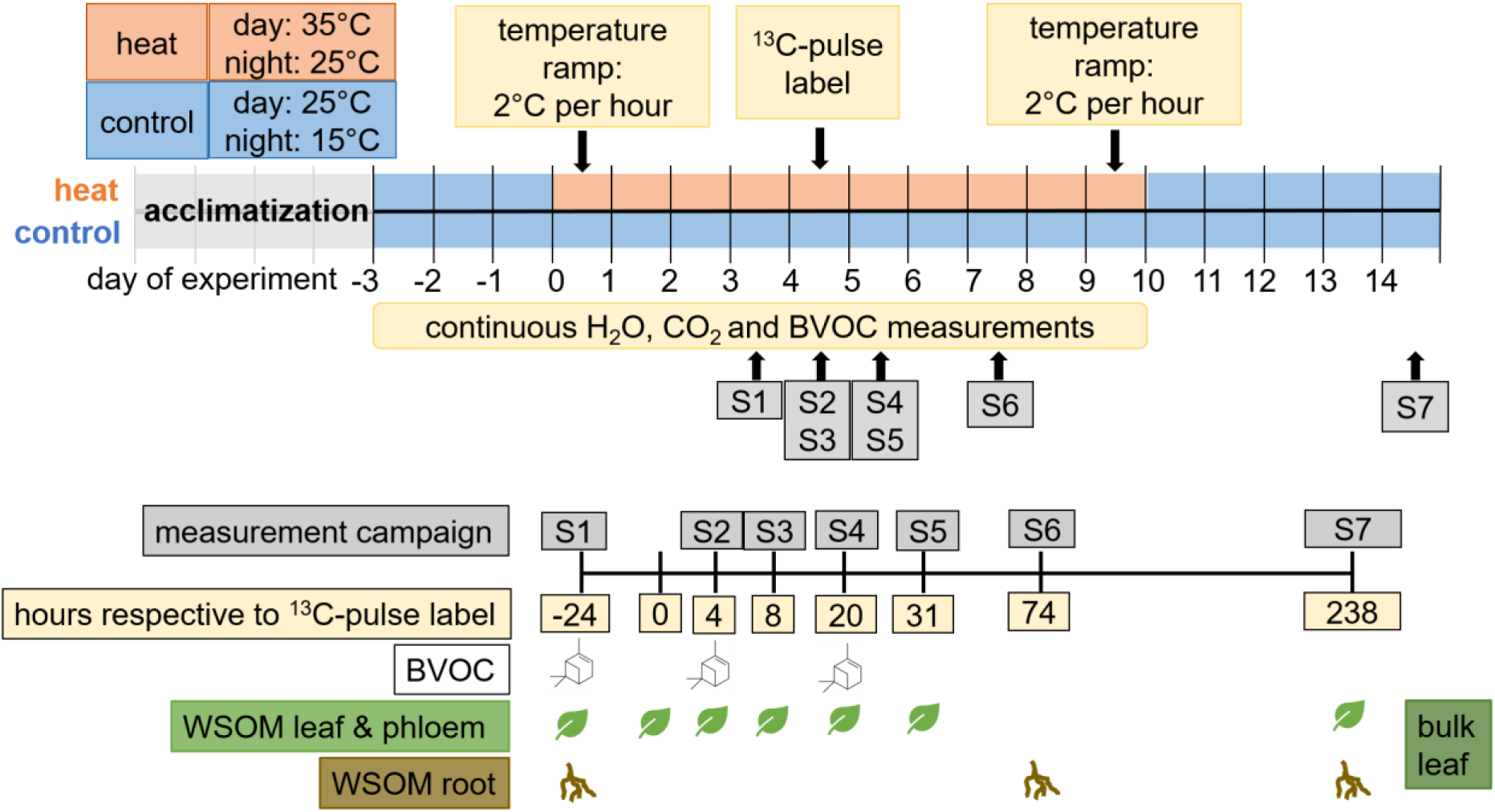
Experimental design showing the timeline of continuous and discrete measurements as well as the timing of heat stress. Biogenic volatile organic compound emissions (VOC) were measured continuously via a PTR-ToF-MS and campaign-wise with adsorbent tubes filled with Tenax. Water soluble organic matter (WSOM) was sampled from phloem of branchlets, leaf tissue and roots.

### Measurements of biogenic volatile organic compounds

The proton-transfer-reaction time-of-flight mass spectrometer (PTR-ToF-MS 4000, Ionicon Analytic, Innsbruck, Austria) for the continuous measurements of VOC emissions was operated at 80°C, 2.7 mbar drift pressure and 500 V drift voltage producing an E/N of 128 Td. A multi-component gas mixture (Apel Riemer Environmental, USA) and a liquid calibration unit (Ionicon Analytic) were used for calibration over different humidity steps as described in Meischner et al. (2024). The measured protonated mass-to-charge ratios (m/z) were analyzed using the Ionicon data analyzer software (IDA, Ionicon Analytic, Innsbruck, Austria) and compounds were assigned via the GLOVOCs data base (Yáñez-Serrano et al., 2021). Emission flux of the identified compounds was then calculated using Eqn. S5 (Supporting Information). For further analysis, only the compounds with emissions larger than the limit of detection, i.e. 3σ of the concentration measured in the empty reference cuvette, were selected (Supporting Information Table S2). For the calculation of carbon allocation to VOC fluxes, the molar emission rates of the compounds were multiplied by the number of C atoms present in the compound (Kesselmeier et al., 2002). Due to the high diversity of emitted compounds, only monoterpene (MT, m/z 137.13) and sesquiterpene (SQT, m/z 205.20) emissions and their respective fragments and isotopes were further analyzed, since they are considered as important heat stress markers (Bourtsoukidis et al., 2024; Duhl et al., 2008; Holopainen et al., 2018, 2025; Vickers et al., 2009).

Additionally, on three measurement points before and after the ^13^CO_2_ pulse label (Figure 1), terpenoid emissions were sampled on thermodesorption tubes filled with 80 mg Tenax (Sigma Aldrich, Munich, Germany) for two hours at a flow rate of 200 ml min^-1^ to identify distinct terpenoids and their respective ẟ^13^C ratios. Adsorbed compounds were analyzed by gas chromatography (GC, 7890B, Agilent Technologies, Böblingen, Germany), see Haberstroh et al. (2019) and Kreuzwieser et al. (2021) for a detailed description. Briefly, samples were heated to 240°C (thermodesorption unit, TDU, Gerstel, Mühlheim a.d. R., Germany) and cryo-focused at -70°C (cold injection system, CIS, Gerstel). The CIS was then heated to 240°C and the analytes released onto a separation column (DB-5MS-UI, Agilent Technologies). 10% of the eluate was channelled into a mass spectrometer (5975C, Agilent) operating at 70eV, with an ion source temperature of 230°C and a quadrupole temperature of 150°C, while 90% of the eluate was directed to a combustion furnace (GC5 Interface, Elementar, Hanau, Germany) operating at 850°C, which was coupled to an isotope ratio mass spectrometer (Isoprime, precisION, Elementar). Analysis of ẟ^13^C relative to the Vienna Pee Dee Belemnite standard (VPDB, Supporting Information Eqn. S7) was done using the lyticOS software (Elementar), while the GC-MS signals were analyzed using the Mass Hunter Software (version B.07.00/7.0.457.0, Agilent Technologies; Haberstroh et al., 2019). A standard mixture of five compounds (α-pinene, β-pinene, limonene, trans-β-ocimene, trans-caryophyllene) was used to quantify concentrations of the compounds. All MTs and SQTs which were not included in the standard mixture were quantified using α-pinene and trans-β-caryophyllene, respectively. Emission rates per leaf area were then calculated using Eqn. S6 (Supporting Information). In total, 16 terpenoids could be identified in *P. menziesii* and five terpenoids in *F. sylvatica* (Supporting Information Table S3).

### Water-soluble organic matter and bulk material for ^13^C analysis

For the analysis of water-soluble organic matter (WSOM), little branchlets were cut during the seven sampling campaigns (Figure 1). After removal of the outer dead bark tissue, samples were separated carefully into phloem, xylem and leaf tissue, frozen immediately in liquid nitrogen and stored at -80°C until further analysis. For the extraction of WSOM, frozen samples were ground on liquid nitrogen and 50 mg of the ground material was placed into 1.5 ml of distilled water, heated for five minutes at 90°C and then centrifuged for five minutes at 14000 x g. 150 µl of the supernatant was transferred into tin capsules (5×9 mm, IVA Analysetechnik, Meerbusch, Germany) and dried at 60°C for 48 hours (Wegener et al., 2010). This step was repeated until at least 0.4 mg of residue remained in the capsule. For the analysis of bulk material, separated samples of leaves, phloem, xylem and roots were dried at 60°C for 48 hours, milled (MM400, Retsch GmbH, Haan, Germany) and 2 mg of *F. sylvatica* and 3 mg of *P. menziesii* samples were weighed into tin capsules (5×9 mm, IVA Analysetechnik). δ^13^C was analyzed on an Elemental Analyzer (EA, Vario Isotope Cube, Elementar) coupled to an Isotope Ratio Mass Spectrometer (IRMS, Isoprime precision, Elementar) as detailed in Werner et al. (2009). Briefly, samples were combusted at 950°C and diluted with Helium at a ratio of 6.5%. Diluted samples were then channeled to the IRMS, ionized and ratio of ^12^C to ^13^C relative to VPDB (ẟ^13^C) was analyzed with a precision of 0.05 ‰ (Supporting Information, Eqn. S7).

### Statistical analysis

Statistical analysis was performed using R (R Core Team, 2026).

After checking for normality of residuals and homogeneity of variance (package “performance”), a log-transformation was applied to MT and SQT emissions. Thereafter, continuous measurements were analyzed with linear mixed effect models using the package “lme4” (Bates et al., 2015; Lüdecke et al., 2021). The day of the experiment, tree species, species combination and treatment were used as fixed effects and the individual sapling as a random effect. Estimated marginal means (package “emmeans”) were used for post-hoc analysis (Lenth, 2023).

The same approach was used to compare compound-specific ẟ^13^C and emission fluxes of terpenoids measured by GC-IRMS. Fixed effects were measurement campaign, tree species, species combination, treatment and the respective compound, while the individual sapling was added as a random effect. ẟ^13^C and emission rates were log-transformed to meet model criteria. In both models, species combination had no significant effect (p > 0.05). Moreover, since biomass harvest revealed no visible competition for rooting space, differences between inter- and intraspecific combinations were not further analyzed.

A first order exponential decay function was used to model the turnover of ^13^C in respiration and leaf WSOM (Eqn. 1).

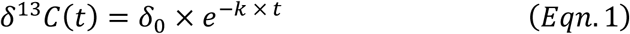

Where ẟ_0_ is the initial ^13^C excess (‰) at t=0, k is the rate constant (day^-1^) and t is the time since the ^13^CO_2_ pulse. The model was fitted per species and treatment using “nlsList” from the “nlme” package (Pinheiro & Bates, 1996). The mean residence time (MRT) and the half-life of the ^13^CO_2_ label were derived from the rate constant (Epron et al., 2012), while standard errors were calculated using error propagation. Significant differences between the two species and treatments were assessed using pairwise z-tests of the rate constant with a Holm correction. Due to the high variability of plant individuals, no model could be fitted for WSOM of phloem and root tissue.

## Results

### Tree biomass and leaf gas exchange

In both species, total biomass was not significantly altered between the heat and control treatment, yet total biomass of *P. menziesii* significantly exceeded that of *F. sylvatica* (p < 0.001, Supporting Information Table S1). However, under heat stress, 69±14% of foliage of *F. sylvatica* turned brown until the end of the experiment with a considerable variation between individuals (2% to 96 %), while foliage of *P. menziesii* remained unaffected.

During all days of heat stress (day 0 to 9 of the experiment), *A_net_* of heat-stressed *P. menziesii* was significantly lower than that of control saplings (1.3±0.1 and 3.2±0.1 µmol m^-2^ s^-1^, respectively, p < 0.05), while *A_net_* of *F. sylvatica* did not differ significantly between the two treatments (p > 0.05, Figure 2 A&B). In both species, *E* was not significantly different between both treatments during the heat stress period due to the high variability between the saplings (Figure 2 C&D). However, compared to the pre-stress period (day -3 to -1 of the experiment), *E* of heat-stressed saplings was significantly increased during the heat stress by 0.32±0.05 mmol m^-2^ s^-1^ in *F. sylvatica* and 0.22±0.03 mmol m^-2^ s^-1^ in *P. menziesii* (p < 0.001). In *P. menziesii*, this response resulted in a significantly lower *WUE* during all days of heat stress compared to control conditions (3.3±0.5 and 9.2±2.1 µmol mmol^-1^, respectively, p < 0.05, Supporting Information Figure S1). In *F. sylvatica, WUE* of heat-stressed saplings was only significantly decreased by 4.9±0.3 µmol mmol^-1^ compared to the respective pre-stress measurements (p < 0.01).

**Figure 2:**
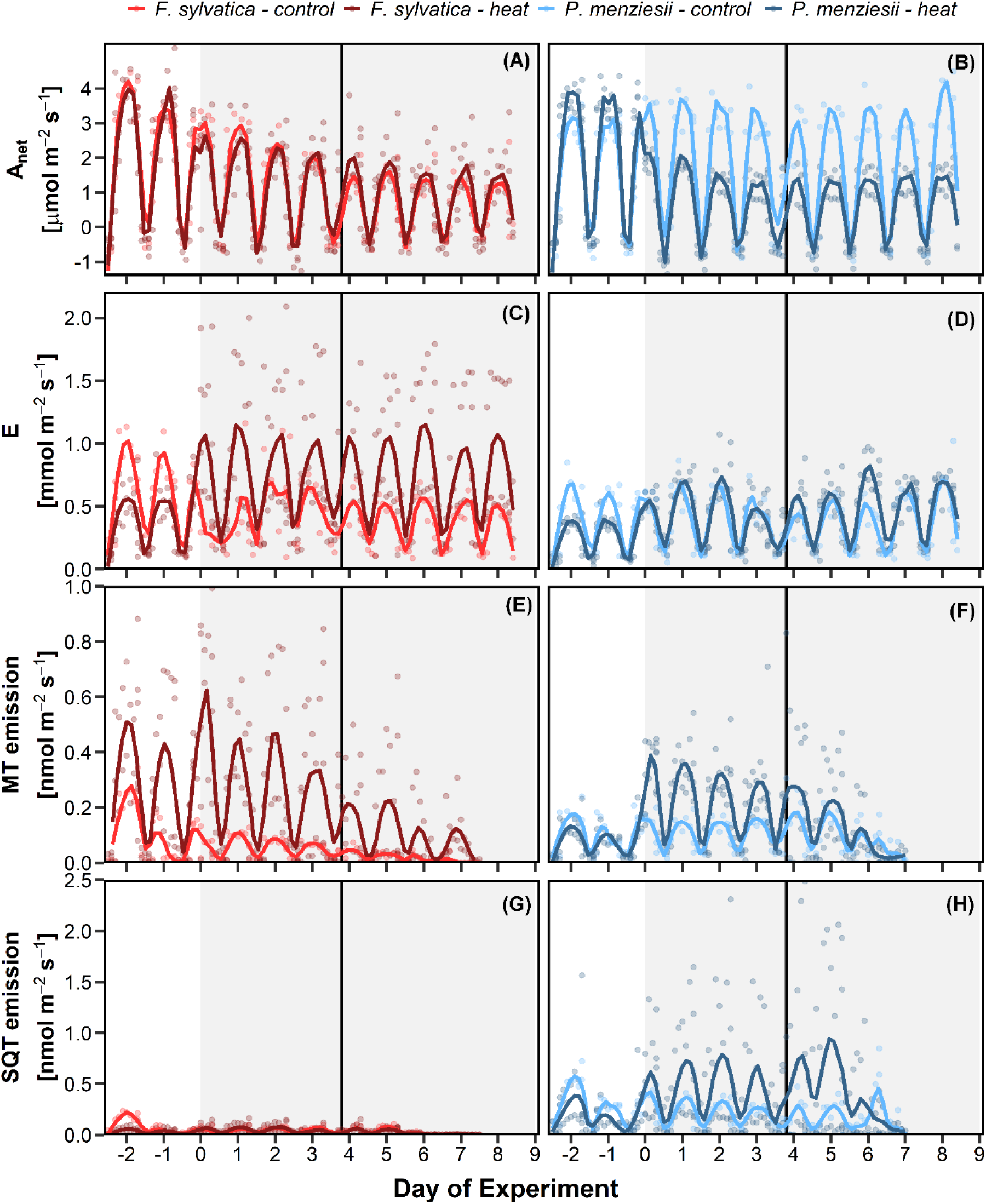
Continuous measurements of net carbon assimilation (A_net_, A&B), transpiration (E, C&D), monoterpene (MT, E&F) and sesquiterpene emissions (SQT, G&H) of *F. sylvatica* (A, C, E, G) and *P. menziesii* (B, D, F, H) under control (25°C, bright color, n=4 per species) and heat stress (35°C, dark color, n=8 per species) conditions. Points show the-hourly mean of all measured individuals, while lines are fitted with a loess smoothing spline with a span of 0.08. Grey shaded area in the background represents the heat stress period, while the black line shows the application of the ^13^CO_2_ pulse label.

### Terpenoid emissions

In both species, heat induced a significant increase in quantity and diversity of emitted terpenoids. Continuous measurements revealed a transient increase of MT emissions during the first three days of heat stress compared to pre-stress measurements, particularly in *P. menziesii* (+0.45±0.07 nmol m^-2^ s^-1^, p < 0.05). However, after six days of heat stress emissions, declined significantly compared to the initial increase by -0.54±0.11 nmol m^-2^ s^-1^ in *F. sylvatica* and -0.48±0.09 nmol m^-2^ s^-1^ in *P. menziesii*, (p < 0.01, Figure 2 E&F). SQT emissions were mainly detected in *P. menziesii*, which tended to be slightly higher during all days of heat stress compared to control conditions (0.62±0.5 and 0.54±0.30 nmol m^-2^ s^-1^, respectively, Figure 2 G&H).

Terpenoid composition was more diverse in *P. menziesii* than in *F. sylvatica* (Figure 3). In *F. sylvatica*, heat stress strongly increased emissions of sabinene and β-phellandrene (210.2 ± 102.2 and 39.2 ± 24.2 nmol m^-2^ h^-1^) compared to control conditions and induced the emission of the acyclic MTs β-myrcene and trans-β-ocimene (41.9 ± 28.2 and 2.2 ± 1.9 nmol m^-2^ h^-1^), which were not detected in the control treatment. Under control conditions, *P. menziesii* mainly emitted (in descending order) farnesene, sabinene and α-pinene comprising 56% of all emitted terpenoids. Under heat stress, highest absolute increase of emission rates (in nmol m^-2^ s^-1^) was detected for the acyclic compounds farnesene (+55.5), β-myrcene (+27.5), β-farnesene (+27.3) and trans-β-ocimene (+23.6) as well as for the cyclic compounds γ-terpinene (+34.7) and sabinene (+31.8). Highest proportional increase in emission rates was found for the compounds with emissions close to zero in the control group (α-terpinene, β-cubenene, β-elemene, γ-terpinene, trans-β-ocimene), which were emitted with 13 to 87 times higher rates in the heat stressed group (Figure 3).

**Figure 3:**
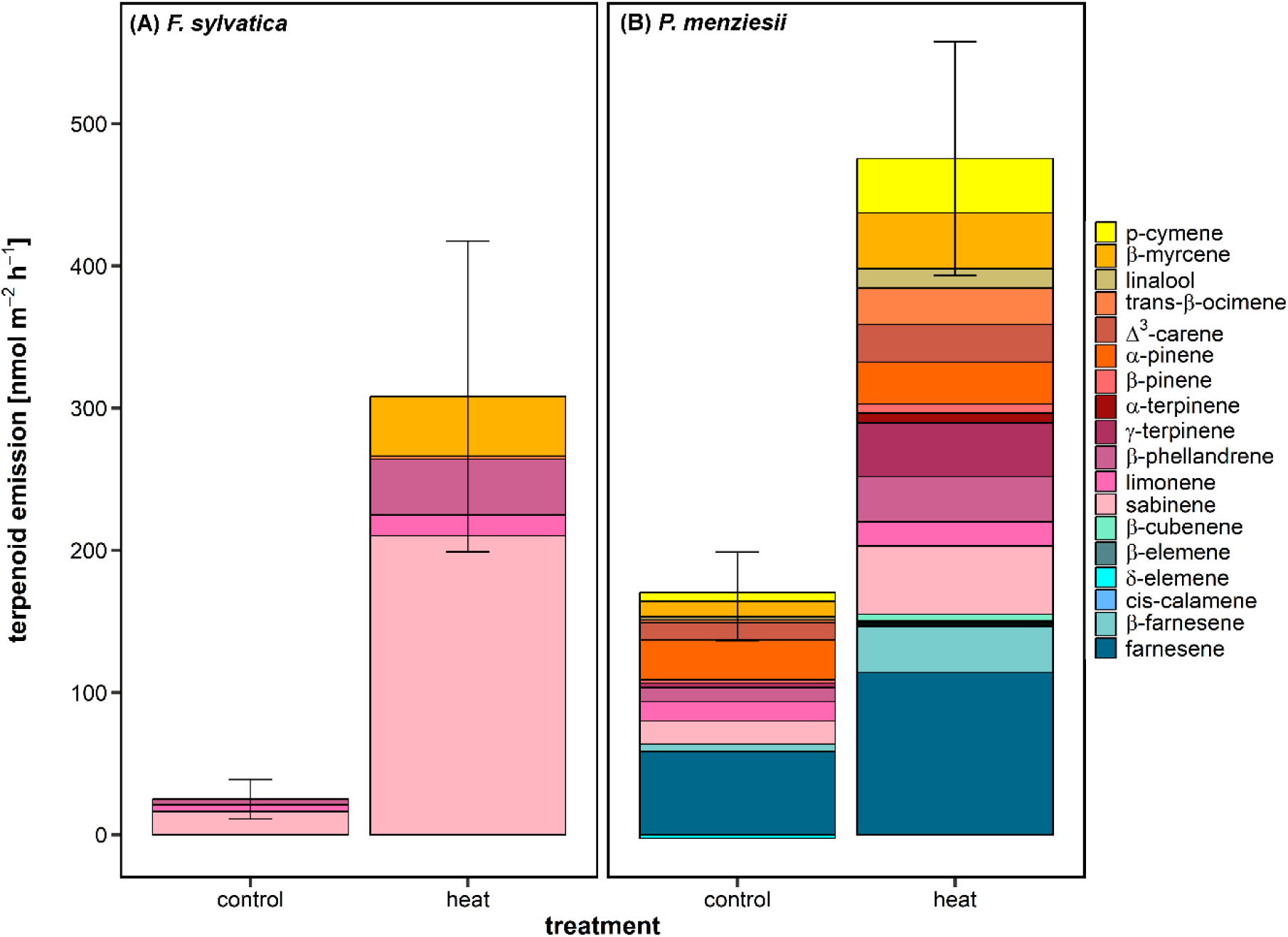
Monoterpene (yellow and reddish colour) and sesquiterpene (green and blue colours) emissions sampled during sampling campaign S1 on day three of the experiment (Figure 1) measured by the GC-MS. Shown are all identified terpene compounds of *F. sylvatica* (A) and *P. menziesii* (B) of the control (n=4) and heat stress (n=8) treatment. Standard error of all compounds was calculated using error propagation.

### Relative contribution of carbon to respiration and BVOC emissions

For both species, heat stress increased relative loss of assimilated carbon via respiratory processes (Fig. 4 A&B). While the increase in respiratory C loss was steady during all days of heat stress in *P. menziesii* (+25.9±3.5 % compared to control)*, F. sylvatica* showed a lower increase during the first three days of heat stress (+11.3±5.3% compared to control) followed by a transient rise of respiratory C loss to up to 52.5±18.8 % of freshly assimilated C during day four to seven of the heat stress period (Figure 4 A&B). Both species steadily increased C allocation into VOCs until the fourth day of heat stress (+1.2 % in *F. sylvatica* and +1.5 % in *P. menziesii*). Concurrent with decreasing MT emissions (Figure 2 E&F), C allocation into VOCs declined after six days of heat stress to 0.2±0.0 % and 0.5±0.2 % of total C uptake in *P. menziesii* and *F. sylvatica*, respectively (Figure 4 C&D).

**Figure 4:**
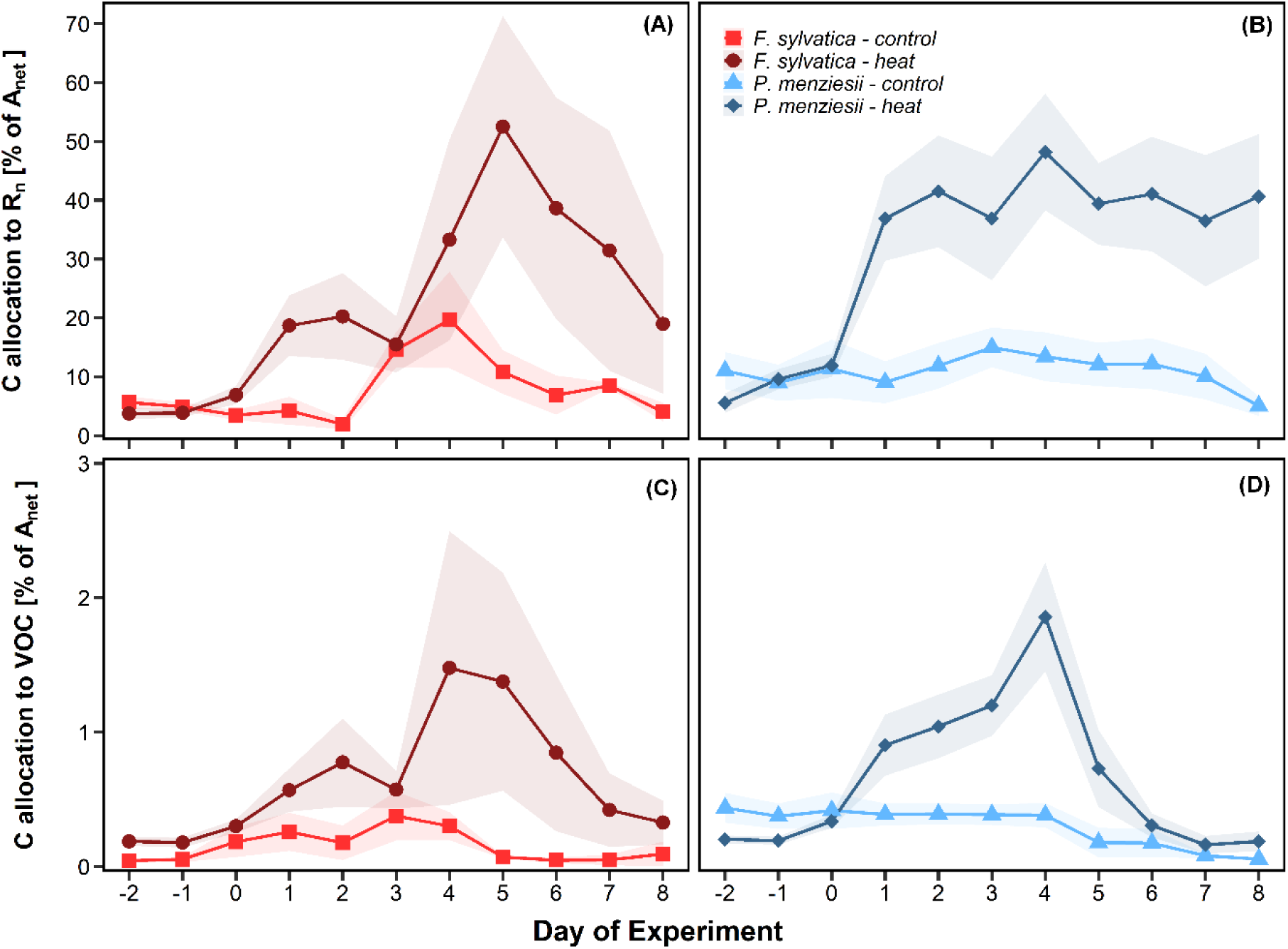
Proportional allocation of total assimilated carbon to nocturnal respiration (*R_n_*, A&B) and to all VOC emissions detected by the PTR-ToF-MS (C&D, Supporting Information Table S1) of *F. sylvatica* (A&C) and *P. menziesii* (B&D). Bright color represents the control (n=4 per species) and dark color the heat stress treatment (n=8 per species). Shaded areas show the standard errors, while points represent the daily mean of all individuals. Calculations are based on the carbon mass of the identified VOCs (Kesselmeier et al., 2002).

### Fate of recently fixed ^13^CO_2_

In both treatments, saplings of *F. sylvatica* assimilated similar amounts of the supplied ^13^CO_2_ (control: 31.6 ± 6.2 %, heat: 29.8 ± 5.2 %), while ^13^CO_2_ uptake of *P. menziesii* was higher in the control (44.8 ± 4.8 %) than in the heat group (32.7 ± 5.5 %). Only a minor proportion of the recently fixed ^13^CO_2_ was detected in water-soluble organic matter (WSOM) of leaves, phloem and roots of both species (Table 1). WSOM of leaves and phloem of *F. sylvatica* was more enriched with ^13^C than that of *P. menziesii*, yet for both species, no significant differences were found between the two treatments. However, MRT of both species responded differently: under control conditions, a lower MRT of ^13^CO_2_ in leaf WSOM was found in *F. sylvatica* (2.5 hours) than in *P. menziesii* (39 hours, Table 2). Under heat stress, this pattern was inverted with a lower MRT of ^13^CO_2_ in leaf WSOM of *P. menziesii* (14 hours) compared to *F. sylvatica* (21 hours).

**Table 1:** ẟ^13^C (‰) of water-soluble organic matter of leaf, phloem and root tissue 24 hours before (S1) and 4 to 238 hours after the ^13^C pulse label (S2-S7).

|  |  | S1<br>pre-label | S2<br>4 hours | S3<br>8 hours | S4<br>20 hours | S5<br>31<br>hours | S7<br>238<br>hours |
| --- | --- | --- | --- | --- | --- | --- | --- |
| *Fagus sylvatica* - control | leaves | -25.8 ± 0.5 | -7.4 ± 13* | -17.8 ± 3.2* | -17.2 ± 4.6* |  | -24.0 ± 1.0 |
|  | phloem | -26.6 ± 0.4 | -20.7 ± 2.4* | -21.4 ± 1.1* | -22.2 ± 1.1 |  | -23.0 ± 1.1 |
|  | roots | -24.4 ± 0.4 |  |  |  | -24.6 ± 1.0 | -23.9 ± 1.3 |
| *Fagus sylvatica* - heat | leaves | -27.5 ± 0.6 | -7.0 ± 6.8* | -11.4 ± 2.8* | -16.9 ± 1.7* |  | -26.2 ± 0.7 |
|  | phloem | -27.2 ± 0.6 | -21.2 ± 0.8* | -22.7 ± 0.9* | -22.7 ± 0.8* |  | -26.0 ± 0.6 |
|  | roots | -25.3 ± 0.3 |  |  |  | -23.6 ± 1.1 | -24.6 ± 0.8 |
| *Pseudotsuga menziesii* - control | leaves | -27.0 ± 0.4 | -19.1 ± 0.5* | -20.9 ± 1.5* | -22.0 ± 0.9* |  | -26.8 ± 0.5 |
|  | phloem | -25.7 ± 0.7 | -25.6 ± 0.6 | -23.8 ± 0.6 | -23.0 ± 0.8 |  | -24.8 ± 0.4 |
|  | roots | -25.5 ± 0.5 |  |  |  | -24.3 ± 0.8 | -24.7 ± 0.4 |
| *Pseudotsuga menziesii* - heat | leaves | -27.1 ± 0.4 | -17.0 ± 1.5* | -20.2 ± 1.6* | -22.9 ± 0.8* |  | -26.1 ± 0.6 |
|  | phloem | -25.9 ± 0.5 | -23.5 ± 0.9 | -23.4 ± 0.7 | -23.6 ± 0.7 |  | -24.9 ± 0.6 |
|  | roots | -25.5 ± 0.6 |  |  |  | -24.4 ± 0.3 | -24.9 ± 0.3 |

Shown are mean values ± standard error of the control (n=4) and heat-stressed (n=8) group.
\* indicate a significant difference compared to pre-stress (S1) measurements, $p < 0.05$ .

**Table 2:**
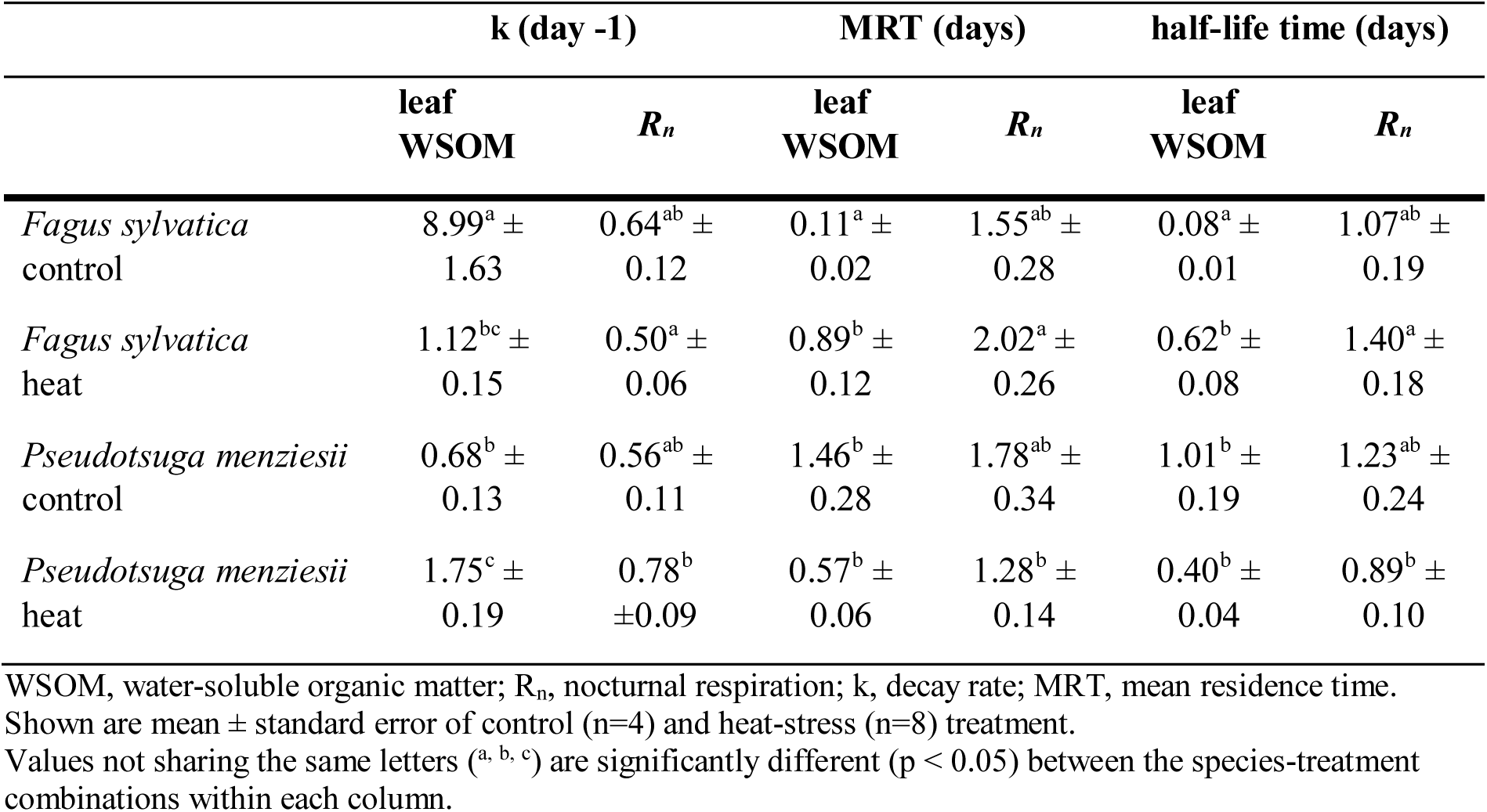
Parameters obtained by the exponential decay function of ^13^CO_2_ in water-soluble organic matter and dark respiration (Eqn. 1).

|  | k (day<sup>-1</sup>) |  | MRT (days) |  | half-life time (days) |  |
| --- | --- | --- | --- | --- | --- | --- |
| | leaf<br>WSOM | $R_n$ | leaf<br>WSOM | $R_n$ | leaf<br>WSOM | $R_n$ |
| *Fagus sylvatica*<br>control | 8.99<sup>a</sup> ±<br>1.63 | 0.64<sup>ab</sup> ±<br>0.12 | 0.11<sup>a</sup> ±<br>0.02 | 1.55<sup>ab</sup> ±<br>0.28 | 0.08<sup>a</sup> ±<br>0.01 | 1.07<sup>ab</sup> ±<br>0.19 |
| *Fagus sylvatica*<br>heat | 1.12<sup>bc</sup> ±<br>0.15 | 0.50<sup>a</sup> ±<br>0.06 | 0.89<sup>b</sup> ±<br>0.12 | 2.02<sup>a</sup> ±<br>0.26 | 0.62<sup>b</sup> ±<br>0.08 | 1.40<sup>a</sup> ±<br>0.18 |
| *Pseudotsuga menziesii*<br>control | 0.68<sup>b</sup> ±<br>0.13 | 0.56<sup>ab</sup> ±<br>0.11 | 1.46<sup>b</sup> ±<br>0.28 | 1.78<sup>ab</sup> ±<br>0.34 | 1.01<sup>b</sup> ±<br>0.19 | 1.23<sup>ab</sup> ±<br>0.24 |
| *Pseudotsuga menziesii*<br>heat | 1.75<sup>c</sup> ±<br>0.19 | 0.78<sup>b</sup><br>±0.09 | 0.57<sup>b</sup> ±<br>0.06 | 1.28<sup>b</sup> ±<br>0.14 | 0.40<sup>b</sup> ±<br>0.04 | 0.89<sup>b</sup> ±<br>0.10 |

WSOM, water-soluble organic matter; $R_n$ , nocturnal respiration; k, decay rate; MRT, mean residence time. Shown are mean ± standard error of control (n=4) and heat-stress (n=8) treatment. Values not sharing the same letters (<sup>a</sup>, <sup>b</sup>, <sup>c</sup>) are significantly different ( $p < 0.05$ ) between the species-treatment combinations within each column.

^13^C enrichment of respired CO_2_ was highest in the night directly after the pulse label and declined exponentially over the following two nights (Figure 5). Similar to the reversed patterns observed in WSOM, we observed a divergent response of respired ^13^CO_2_: under heat stress, faster decay rates were observed in *P. menziesii* than in *F. sylvatica* with a half-life time of 22 and 34 hours, respectively, whereas in the control treatment, a slower decay was found in *P. menziesii* than in *F. sylvatica* with a half-life time of 30 and 26 hours, respectively (Table 2).

**Figure 5:**
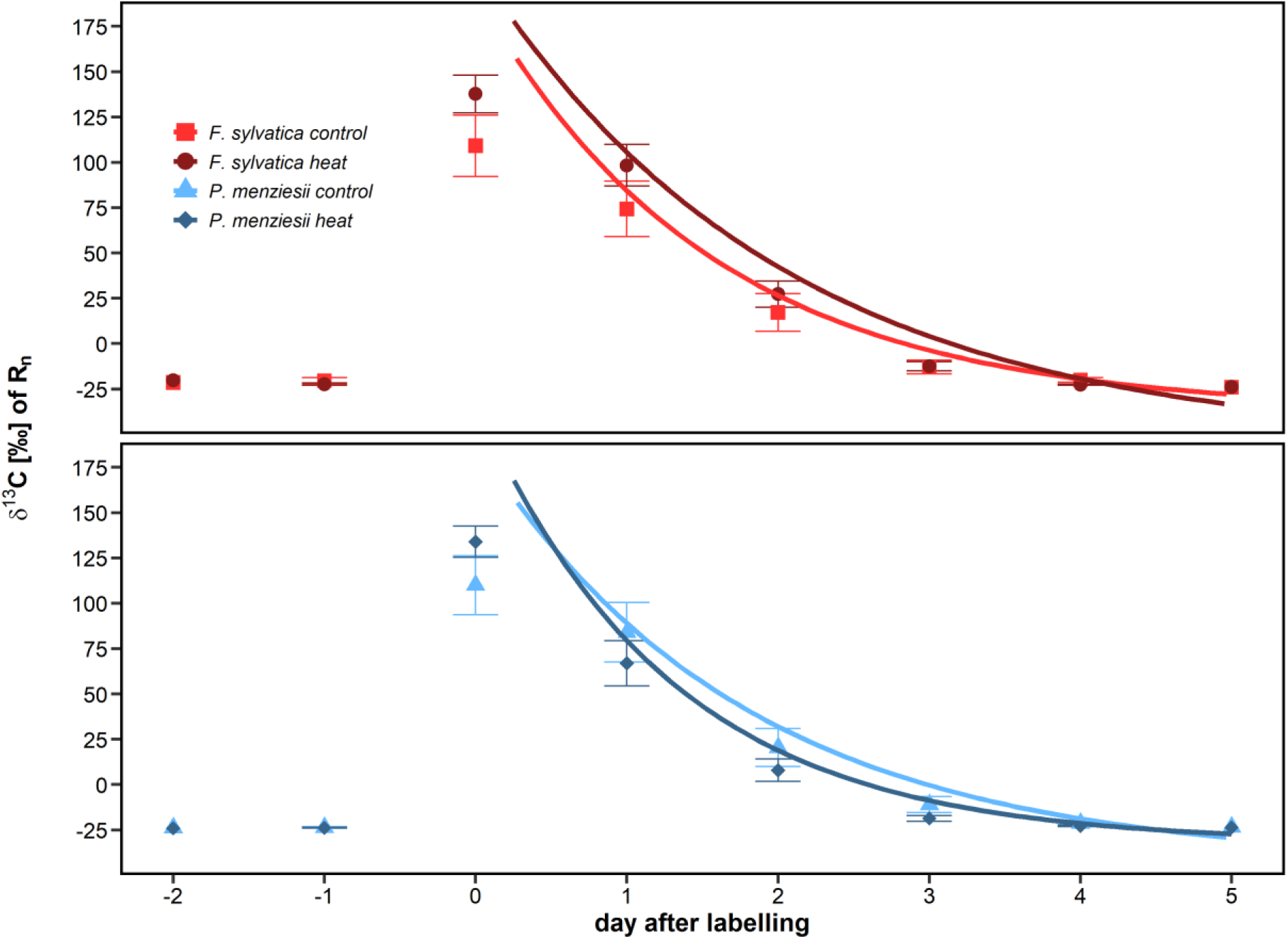
ẟ^13^C (‰) of nocturnal respiration (*R_n_*) of control (n=4 per species) and heat-stressed (n=8 per species) saplings of *F. sylvatica* (A) and *P. menziesii* (B). Points show the mean ẟ^13^C during the seven hour long dark period of all plant individuals, error bars the respective standard error and lines the exponential decay function fitted by Eqn. 1.

After the ^13^C pulse labelling, all emitted MTs of heat stressed *F. sylvatica* were significantly enriched (p < 0.01), whereas under control conditions only sabinene and limonene were slightly enriched 4 and 20 hours after the label, respectively (p < 0.001). MTs with higher emissions under heat stress compared to control (β-myrcene, β-phellandrene, sabinene) also showed the highest enrichment directly after the ^13^C labelling (Figure 6) with a stronger enrichment being directly related to a stronger increase in emissions (Figure 6). However, 20 hours after the pulse label, ẟ^13^C of MTs declined quickly and only sabinene remained significantly enriched (72.3 ± 19.3 ‰, p < 0.001, Figure 6D). A more gradual increase of ẟ^13^C was observed in limonene with highest enrichment 20 hours after the label in the heat (45.2±30.2 ‰) and control treatment (7.4±35.1 ‰, Figure 6 C&G). Emission rates of this compound were only slightly increased under heat (+6.7±2.7 nmol m^-2^ s^-1^). Low emission rates of trans-β-ocimene prevented ẟ^13^C analysis in the IRMS.

**Figure 6:**
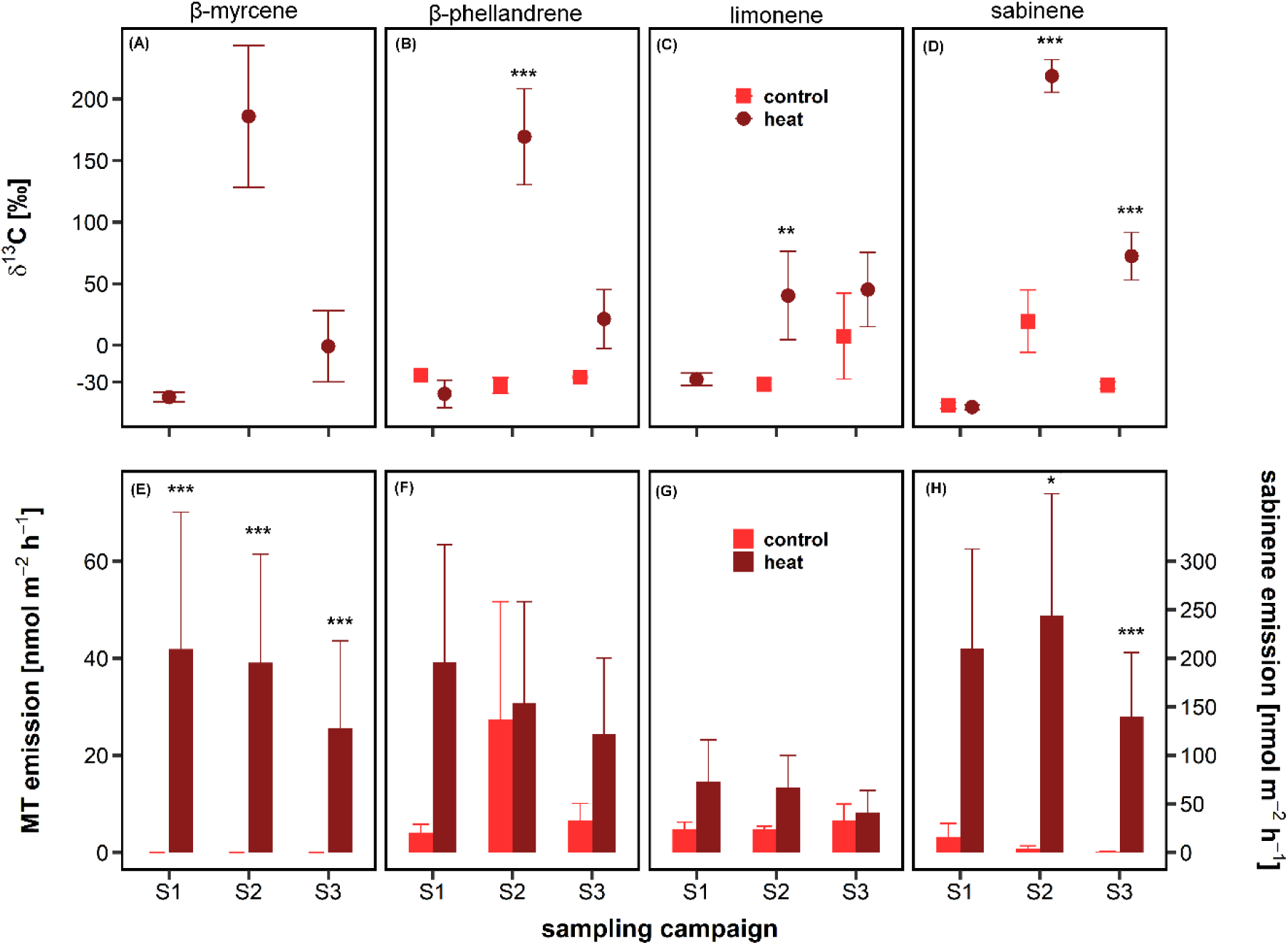
ẟ^13^C (A-D) and respective emission rates (E-H) of monoterpenes emitted by *F. sylvatica* under control (n=4) and heat (n=8) conditions. Shown are mean values and respective standard errors of all individuals. Note that sabinene emission rates in panel H are displayed on a separate scale. Asterisks indicate a significant difference between control and heat stressed saplings (* p < 0.05; ** p < 0.01; *** p < 0.001). Missing points in panel A and C and missing compounds compared to Figure 3 are due to low emission rates, which prevented peak detection in the IRMS. Samples were taken 24 hours prior to (S1) as well as four (S2) and 20 hours (S3) after the ^13^C pulse label (Figure 1).

MT emissions of *P. menziesii* were less ^13^C enriched than in *F. sylvatica* and showed four different patterns (Figure 7): α- and β-pinene had higher emission rates in the control treatment and were not enriched after the pulse label in both treatments. Linalool and β-phellandrene showed a gradual ^13^C enrichment with a stronger increase of ẟ^13^C in the control than in the heat treatment (+20.8 ‰ and +4.8 ‰ compared to pre-label ẟ^13^C, respectively), despite higher emission rates in the heat treatment. β-myrcene, limonene and sabinene showed a more immediate increase (S2) and subsequent decrease (S3) in ẟ^13^C with a similar enrichment in the control and heat treatment. Trans-β-ocimene and γ-terpinene had significantly elevated emission rates (p < 0.05) and a higher ^13^C enrichment in the heat compared to the control treatment. For both compounds, a pronounced variability of δ^13^C between the tree saplings was observed in the control treatment. Three MTs (linalool, β-phellandrene, limonene) tended to be already more enriched in the heat treatment than in the control treatment before the label application (Figure 7 B, E, F).

**Figure 7:**
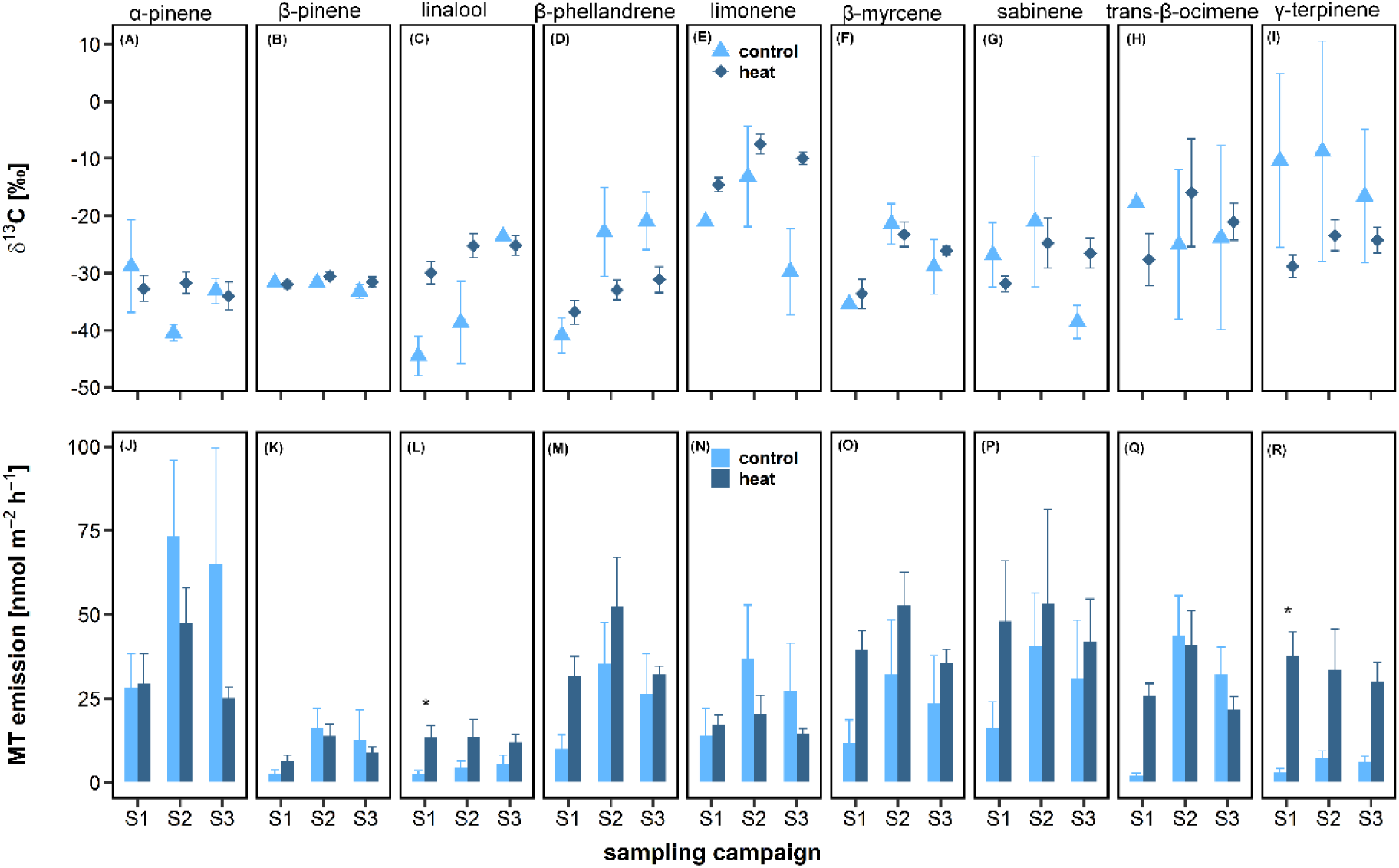
ẟ^13^C (A-I) and respective emissions rates (J-R) of monoterpenes emitted by *P. menziesii* under control (n=4) and heat (n=8) conditions. Shown are mean values and standard errors of all individuals. Asterisks indicate a significant difference between control and heat stressed saplings (* p < 0.05; ** p < 0.01; *** p < 0.001). Missing compounds compared to Figure 3 are due to low emission rates, which prevented peak detection in the IRMS. Samples were taken 24 hours prior to (S1) as well as four (S2) and 20 hours (S3) after the ^13^C pulse label (Figure 1).

The same four patterns were also visible in SQT emissions (Figure 8): no enrichment and lowest emission rates of all terpenoids was detected for β-elemene. δ^13^C of β-cubebene increased more strongly in the control than in the heat treatment (p < 0.01), despite significantly elevated emission rates in the heat group (p < 0.05). β-farnesene was more enriched in the heat compared to the control treatment (-8.9±3.9 ‰ and -21.7±4.3 ‰) and tended to have higher emission rates in the heat group. Farnesene was the most emitted and the most ^13^C enriched terpenoid with an immediate increase (S2) and subsequent decrease (S3) of δ^13^C, which was more pronounced in the control (206.5±74.2 ‰) than in the heat (41.1±24.1 ‰) treatment (p < 0.001). Low emission rates of cis-calamene, α-terpinene and Δ^3^-carene prevented peak detection in the IRMS. Four terpenoids (trans-β-ocimene, α-pinene, β-farnesene, farnesene) responded with a pronounced emission burst to the expansion and movement of the enclosures for the ^13^CO_2_ pulse labelling (Figure 7 L&P, Figure 8 G&H).

**Figure 8:**
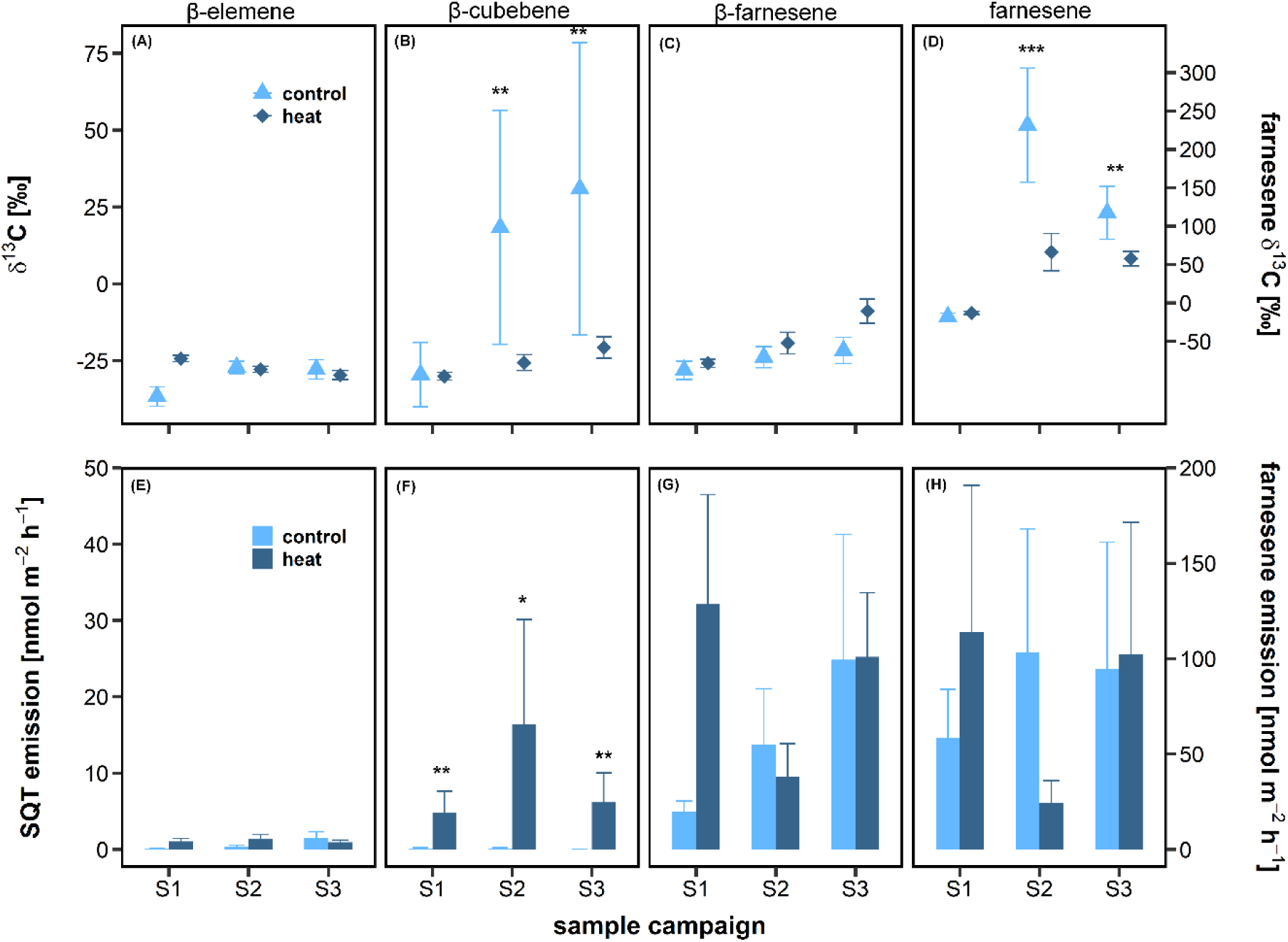
ẟ^13^C (A-D) and respective emissions rates (E-H) of sesquiterpenes emitted by *P. menziesii* under control (n=4) and heat (n=8) conditions. Points and bars show the mean and error bars the respective standard error of all individuals. Asterisks indicate a significant difference between control and heat stressed saplings (* p < 0.05; ** p < 0.01; *** p < 0.001). Note that ẟ^13^C ratios and emission rates of farnesene (D&H) are displayed on a separate scale. Missing compounds compared to Figure 3 are due to low emission rates, which were not sufficient for peak detection in the IRMS. Samples were taken 24 hours prior to (S1) as well as four (S2) and 20 hours (S3) after the ^13^C pulse label (Figure 1).

## Discussion

Globally, forests are increasingly suffering from heat waves constraining ecosystem functioning (Novick et al., 2024; Werner et al., 2025; Yuan et al., 2025). We show that *F. sylvatica* and *P. menziesii* saplings respond to heat stress by prioritizing carbon allocation towards maintenance respiration and protection via VOCs. In *F. sylvatica*, heat stress delayed the export of fresh assimilates, whereas in *P. menziesii,* turnover of fresh assimilates was accelerated. Analysis of compound-specific δ^13^C showed that *F. sylvatica* increased *de novo* synthesis of specific terpenoids, altering both terpenoid composition and emissions. In contrast, *P. menziesii* mainly released stored terpenoids, albeit elevated *de novo* synthesis of specific compounds suggested a selective upregulation for thermal protection.

### Heat stress significantly alters carbon allocation and water use efficiency

Under heat stress, both species prioritized allocation of recently fixed ^13^CO_2_ to maintenance respiration and VOC emissions over allocation to WSOM (Figure 4, Table 1). WSOM serves as an immediate C pool providing substrate for new growth, respiration, defence and transport, and pools are generally kept above a species-specific minimum (Hartmann & Trumbore, 2016; Huang et al., 2021; Martínez-Vilalta et al., 2016). In *Picea abies*, C deprivation led to preferential allocation of fresh assimilates to maintenance respiration and terpenoids at the expense of other C sinks (Huang et al., 2018, 2021). Moreover, simulated herbivory induced *de novo* synthesis of terpenoids in *P. abies*, which was fuelled by the mobilization of locally stored WSOM and starch (Huang et al., 2020). We see three reasons for the low ^13^C incorporation into WSOM in our study: (1) strength of our pulse label (283‰) was moderate (e.g. Rehschuh et al., 2022; Ruehr et al., 2009; Zang et al., 2014), (2) existing WSOM pools diluted the labelling signal (Keel et al., 2006, 2012), and (3) WSOM was directly re-invested into *de novo* synthesis of VOCs and maintenance respiration.

Interestingly, carbon pool turnover diverged between the two species: accelerated C turnover was found in *P. menziesii* (Table 2), which was probably stimulated by the high respiratory cost for e.g. membrane and enzyme stabilization (Scafaro et al., 2021; Teskey et al., 2015). Contrarily, heat reduced C turnover rates in *F. sylvatica*, which was also found by Blessing et al. (2015), but is in contrast to Dannoura et al. (2011). Additionally, carbon investment into maintenance respiration and terpenoids decreased after six days of heat stress (Table 2, Figure 4). Thus, reduced turnover was likely driven by decreased net assimilation and inhibited sucrose synthesis, which can decrease WSOM pools (Scafaro et al., 2021) and ultimately limit available substrate for both respiration (Bruhn et al., 2022; Jones et al., 2024) and terpenoid synthesis (Huang et al., 2020). Such a strategy may ensure that soluble sugars do not drop below a safety margin to prevent excessive C exhaustion (Huang et al., 2018, 2021), particularly since maintenance respiration already consumed up to 50% of fresh assimilates clearly reducing net carbon gain. For the future, a potential shift towards higher carbon loss relative to carbon uptake is expected (Niu et al., 2024), even though thermal acclimation of photosynthesis might alleviate increased carbon loss via maintenance respiration (Deluigi et al., 2025; Xiaoni et al., 2025).

Reduced net carbon uptake coincided with increased transpiration resulting in lower water use efficiency, which can prevent overheating of leaves under heat stress (Marchin et al., 2023). Such stomatal decoupling occurs under both unlimited water supply (Diao et al., 2024; Drake et al., 2018) and soil drought (Marchin et al., 2023), albeit severe edaphic drought ultimately impeded evaporative cooling (Rehschuh et al., 2022; Urban et al., 2017) and increased leaf mortality (Marchin et al., 2023; Zeppan et al., 2026). However, even under ample water supply, increased transpiration may quickly deplete soil water reserves (Diao et al., 2024; Urban et al., 2017) potentially constraining the protective effect under long-term heat extremes.

### δ^13^C reveals compound-specific function of terpenoids for protection against heat

Beyond increased maintenance respiration and evaporative cooling, chemical protection via the synthesis of specific terpenoids provides an additional protective mechanism against thermal stress (Vickers et al., 2009). Particularly in heat-stressed saplings of *F. sylvatica*, some terpenoids (sabinene, β-myrcene, β-phellandrene) showed a strong incorporation of ^13^C indicating high *de novo* production. Most broadleaved species, such as *F. sylvatica*, do not possess specific storage tissues for VOCs, thus production depends on light, temperature and substrate availability (Holopainen et al., 2025; Kleist et al., 2012). Increasing *T_air_* (up to ∼43°C) generally led to increased MT emissions in *F. sylvatica* (Dindorf et al., 2006; Holzke et al., 2006; Llusia et al., 2013; Šimpraga et al., 2011), yet in several tree species reduced assimilation rates under stress limited substrate availability and decreased MT emissions (Dindorf et al., 2006; Niinemets et al., 2010; Staudt et al., 2002; Staudt & Bertin, 1998; Werner et al., 2020; Yáñez-Serrano et al., 2019). This might explain our finding that after an initial increase, MT emissions decreased after around six days of heat stress MTs have been shown to increase thermotolerance of membranes and reduce oxidative cell damage by ROS enabling higher assimilation rates under heat stress than in non-emitting species (Delfine et al., 2000; Loreto et al., 1998; Peñuelas & Llusià, 2002; Vickers et al., 2009). Particularly, an increased production of acyclic compounds was observed and explained by a potential reduction of enzymatic activity, which possibly impedes cyclization of the precursor geranyl diphosphate (Jardine et al., 2017; Staudt & Bertin, 1998). We did indeed find increased *de novo* synthesis of acyclic β-myrcene and trans-β-ocimene under heat stress, but also of bicyclic sabinene and cyclic β-phellandrene. Most likely, 35°C were not sufficient to elucidate a shift from cyclic to acyclic MT emissions as observed in previous studies with *Quercus ilex* at 43-55°C (Loreto et al., 1998; Staudt & Bertin, 1998), but we still conclude that acyclic MTs seemingly play an important role for thermal protection.

In *P. menziesii*, we also found increased emissions of acyclic MTs (β-myrcene, linalool, trans-β-ocimene), but also acyclic SQTs (β-farnesene, farnesene) under heat stress (Figure 3). Particularly the acyclic SQTs were strongly ^13^C enriched indicating emissions from *de novo* synthesis rather than storage release of those compounds, which was also found for α- and β-farnesene emissions of drought-stressed *Pinus sylvestris* (Kreuzwieser et al., 2021) and linalool and SQT emissions of light limited *P. abies* (Huang et al., 2020). In general, ^13^C enrichment was less pronounced in *P. menziesii* compared to *F. sylvatica* indicating that emissions were dominated by stored terpenoids rather than by *de novo* synthesized compounds. Previous studies showed a non-negligible contribution of *de novo* synthesis to terpenoid emissions in conifers accounting for 10 to 39% of total emissions in *P. sylvestris* (Ghirardo et al., 2010; Kleist et al., 2012), 7% in *Larix decidua* and 23% in *P. abies* (Ghirardo et al., 2010), 74% in *Pinus ponderosa* (Harley et al., 2014) and 80% in *Pinus halepensis* (Staudt et al., 2017). It has to be noted that cuvette enlargement during the ^13^CO_2_ pulse labelling might have slightly biased our results, since mechanistic stress can induce emissions of stress-related terpenoids (Duhl et al., 2008; Meischner et al., 2025), but was unavoidable due to logistic reasons.

Selected compounds (β-myrcene, limonene, sabinene, farnesene) showed immediate incorporation of ^13^C in both treatments suggesting that fresh assimilates were involved in the *de novo* synthesis. Yet, for trans-β-ocimene and γ-terpinene, this pattern was only found under heat stress together with strongly elevated emission rates (Figure 7). Strong *de novo* synthesis of ocimenes under heat stress was found in various studies (Jardine et al., 2017; Ladd et al., 2023; Nagalingam et al., 2023; Staudt et al., 2017; Staudt & Bertin, 1998), while high ^13^C enrichment was found for γ-terpinene in *P. ponderosa* (Harley et al., 2014), for sabinene and myrcene in *P. abies* (Daber et al., 2025) and for myrcene, α- and β-farnesene in *P. sylvestris* (Kreuzwieser et al., 2021). Interestingly, trans-β-ocimene, β-myrcene and farnesene are acyclic compounds, as discussed above, while γ-terpinene and sabinene share the same synthesis pathway being produced from the terpinen-4-yl cation after a hydride shift of the α-terpinyl-cation (Degenhardt et al., 2009). Since *de novo* synthesis of sabinene in *F. sylvatica* was also strongly increased under heat stress, we conclude that strategic investment into both acyclic compounds and compounds derived from the terpinen-4-yl precursor might enhance thermotolerance of plants.

Other MTs and SQTs (linalool, β-phellandrene, β-cubebene, β-farnesene) of *P. menziesii* were enriched more gradually similarly to limonene emissions of *F. sylvatica.* Several studies found substantial contributions of cytosolic pyruvate as an alternative carbon source for terpenoid production, if fresh assimilates are limited under heat or drought stress (Daber et al., 2025; Ladd et al., 2023; Werner et al., 2020; Yáñez-Serrano et al., 2018). Cytosolic pyruvate is a direct product of starch hydrolysis which can be transported into the chloroplast for terpenoid synthesis in the MEP pathway or converted into Acetyl-CoA for cytosolic synthesis via the MVA pathway (Ladd et al., 2023; Werner et al., 2020; Yáñez-Serrano et al., 2019). For instance in *P. abies*, cytosolic pyruvate was a strong source for limonene synthesis (Daber et al., 2025), and in several Mediterranean species limonene emissions were light-independent, thus did not directly rely on fresh assimilates (Llusià & Peñuelas, 2000; Staudt et al., 1997, 2000). In *Pinus edulis,* terpenoid synthesis was found to be correlated with soluble sugars derived from starch hydrolysis (Trowbridge et al., 2021), while in *P. abies*, MT emissions remained stable under C deprivation at the expense of strongly depleted starch reserves (Huang et al., 2018). Thus, we speculate that partially terpenoid synthesis was fueled by an alternative carbon source, such as intermediately stored starch, which delayed enrichment of those compounds.

Bicyclic α-& β-pinene were not enriched with ^13^C indicating negligible *de novo* synthesis of those compounds. Pinenes and camphene were found to be the most abundantly stored compounds in *P. menziesii* (Duan et al., 2019; Kleiber et al., 2017) and generally show stable emissions without a strong response to changes in environmental conditions (Daber et al., 2025; Hakola et al., 2017; Helmig et al., 2013). In summary, our results suggest that the benefits of enhanced stress defense appear to offset the metabolic cost of selected terpenoids under heat stress, despite the already enhanced carbon loss via maintenance respiration. For future studies, a targeted investigation of the trade-off between carbon allocation into NSCs, secondary metabolites and growth will crucially contribute to our understanding of carbon dynamics in ecosystems under stress (Hartmann & Trumbore, 2016; Huang et al., 2018, 2019, 2024).

### Implications for forest ecosystems under heat stress

In line with our results, there is growing evidence that temperate ecosystems have considerably reduced net carbon uptake, particularly after compound droughts such as in 2018 and 2022 (Thompson et al., 2020; van der Woude et al., 2023). While in most ecosystems annual carbon fixation still exceeds respiration (Pan et al., 2024; Pohl et al., 2023; Scapucci et al., 2024), some extreme examples have already turned into almost permanent carbon sources (Carle et al., 2025; Haberstroh et al., 2022, 2025). The increasing number and intensity of heat waves in Central Europe (Beobide-Arsuaga et al., 2025) will most likely accelerate carbon loss due to the need for protective mechanisms. Albeit the absolute C losses via respiration exceed those via VOC emissions, increasing emissions might have a strong impact on the formation and properties of secondary organic aerosols, particularly if emissions are dominated by acyclic compounds (Faiola et al., 2019; Liu et al., 2026; Mentel et al., 2013). The combination of heat with edaphic drought, where VOC emissions initially increase, but thereafter decline might lead to synergistic effects (Bonn et al., 2019; Daber et al., 2025; Haberstroh et al., 2018; Staudt et al., 2002; Werner et al., 2021; Wu et al., 2015). Such synergies will potentially complicate emission models and predicted impacts on atmospheric chemistry under future climate warming (Guenther et al., 2012). Thus, whether overall VOC emissions from ecosystems will increase or decrease, will depend on the timing and duration of heat waves and edaphic drought in the future.

In summary, we demonstrate that heat stress alters carbon pool turnover and shifts carbon allocation towards maintenance respiration and the production of terpenoids for thermal protection. Our findings suggest that under future temperature extremes, the increased metabolic cost of thermotolerance, particularly through *de novo* synthesis of protective terpenoids and maintenance respiration, may significantly diminish net carbon uptake of temperate forests.

## Acknowledgements

We gratefully acknowledge financial support by the German Research Foundation via the CRC1537 (ECOSENSE, Project ID: 459819582) and the RU Forest Floor (WE 2681/13-1), and by the Studienstiftung des deutschen Volkes. We thank Alexandra Paul and Anne-Marie Schiphorst for assistance with IRMS analysis, Monika Eiblmeier for assistance with GC-IRMS analysis and Eva Schottmüller, Finn Reimold and Lennart Nettler for support with the plant material.

## Author contributions

Study design: CW, SH

Experiment: SD, CS, MM, PLM, HV, MW, KK

Data collection: SD, CS, MM, PLM, HV, MW, KK

Data analysis: SD, CS, MM, PLM, HV, JK, SH

Data interpretation: SD, CS, MM, CW, SH

Manuscript writing: SD, CS and SH with inputs from all authors

All authors critically reviewed the manuscript.

## Data availability

The data that support the findings of this study are available from the corresponding author upon reasonable request.

