## Supporting Information for "^13^CO_2_ pulse labelling reveals species-specific alterations in carbon allocation and volatile organic compound emissions under heat stress"

**Supporting tables**

Table S 1: Dry weight [g] of the different tree tissues and root to shoot ratio [%]. Shown is the mean value of all saplings per treatment (control: n=4 and heat: n=8) and the respective standard error.

|  | *F. sylvatica* control | *F. sylvatica* heat | *P. menziesii* control | *P. menziesii* heat |
| --- | --- | --- | --- | --- |
| total tree | 67.54 ± 19.04 | 55.19 ± 6.20 | 109.36 ± 16.53 | 132.78 ± 26.78 |
| foliage | 6.12 ± 1.98 | 5.12 ± 1.02 | 27.44 ± 6.62 | 33.67 ± 7.19 |
| twig and stem phloem | 10.35 ± 2.22 | 9.40 ± 1.17 | 20.28 ± 1.83 | 25.17 ± 4.35 |
| twig and stem xylem | 32.30 ± 8.07 | 25.42 ± 2.38 | 30.48 ± 4.79 | 33.96 ± 4.42 |
| roots | 18.78 ± 7.75 | 15.26 ± 2.57 | 31.15 ± 5.79 | 39.99 ± 11.40 |
| root:shoot ratio | 0.38 ± 0.09 | 0.38 ± 0.05 | 0.40 ± 0.04 | 0.38 ± 0.06 |

Table S 2: Identified compounds of VOC emissions measured by the PTR-ToF-MS during the experiment. Compounds were identified using the GLOVOCs data base (Yáñez-Serrano et al., 2021) and the IDA Software (Ionicon Analytics, Innsbruck, Austria). Shown are the protonated mass-to-charge ratios (m/z) and the trivial name of the identified compounds.

| **m/z** | **identified compound** |
| --- | --- |
| 33.03 | Methanol |
| 45.03 | Acetaldehyde |
| 47.04 | Ethanol |
| 59.04 | Acetone |
| 60.04 | Acetone ^13^C labelled |
| 61.02 | Acetic Acid |
| 62.02 | Acetic Acid ^13^C labelled |
| 67.05 | Monoterpene fragment |
| 69.06 | Isoprene |
| 70.06 | Isoprene ^13^C labelled |
| 71.04 | Methyl Vinyl Ketone |
| 72.04 | Methyl Vinyl Ketone ^13^C labelled |
| 81.06 | Monoterpene/Sesquiterpene fragment |
| 82.06 | Monoterpene/Sesquiterpene fragment ^13^C labelled |
| 99.07 | 2-hexenal/3-hexenal |
| 107.08 | Monoterpene fragment |
| 135.11 | p-cymene |
| 136.11 | Amphetamine |
| 137.13 | Monoterpene |
| 138.13 | Monoterpene ^13^C labelled |
| 149.10 | Estragole/anethole |
| 153.06 | Methyl Salicylate |
| 154.06 | Methyl Salicylate ^13^C labelled |
| 205.20 | Sesquiterpene |

Table S 3: Identified terpenoids measured by the GC-MS during three measurement campaigns where terpenoids were sampled onto thermodesorption tubes.

| **Compound** | **Species** | **Match Factor** | **CAS** | **Retention time** |
| --- | --- | --- | --- | --- |
| α-pinene | *P. menziesii* | 99.15 | 80-56-8 | 20.84 |
| sabinene | *P. menziesii* | 97.51 | 3387-41-5 | 23.66 |
| β-pinene | *P. menziesii* | 98.96 | 127-91-3 | 23.97 |
| β-myrcene | *P. menziesii* | 96.06 | 123-35-3 | 24.67 |
| para-cymene | *P. menziesii* | 95.63 | 99-87-6 | 27.04 |
| limonene | *P. menziesii* | 98.30 | 464-17-5 | 27.16 |
| β-phellandrene | *P. menziesii* | 92.86 | 555-10-2 | 27.28 |
| γ-terpinene | *P. menziesii* | 97.55 | 99-85-4 | 28.78 |
| α-terpinene | *P. menziesii* | 89.94 | 99-86-5 | 30.21 |
| linalool | *P. menziesii* | 96.04 | 78-70-6 | 30.86 |
| δ-elemene | *P. menziesii* | 67.14 | 20307-84-0 | 39.24 |
| β-elemene | *P. menziesii* | 83.83 | 515-13-9 | 40.50 |
| β-farnesene | *P. menziesii* | 83.95 | 18794-84-8 | 41.56 |
| farnesene | *P. menziesii* | 95.20 | 502-61-4 | 42.48 |
| β-cubenene | *P. menziesii* | 80.69 | 13744-15-5 | 42.88 |
| cis-calamene | *P. menziesii* | 70.79 | 483-77-2 | 43.03 |
| sabinene | *F. sylvatica* | 98.75 | 3387-41-5 | 23.52 |
| β-myrcene | *F. sylvatica* | 96.55 | 123-35-3 | 24.56 |
| limonene | *F. sylvatica* | 95.90 | 138-86-3 | 27.05 |
| β-phellandrene | *F. sylvatica* | 91.08 | 555-10-2 | 27.16 |
| trans-β-ocimene | *F. sylvatica* | 84.19 | 3779-61-1 | 28.11 |

Table S 4: ẟ^13^C ratios of bulk material of the different tissues of the saplings sampled at the end of the experiment (sampling campaign S7).

|  | ***F. sylvatica* control** | ***F. sylvatica* heat** | ***P. menziesii* control** | ***P. menziesii* heat** |
| --- | --- | --- | --- | --- |
| **leaves** | -25.8 ± 0.2 ‰ | -26.6 ± 0.6 ‰ | -27.9 ± 0.3 ‰ | -27.2 ± 0.3 ‰ |
| **phloem** | -25.8 ± 0.7 ‰ | -26.5 ± 0.7 ‰ | -25.5 ± 0.6 ‰ | -26.1 ± 0.2 ‰ |
| **xylem** | -24.2 ± 0.8 ‰ | -25.4 ± 0.5 ‰ | -25.3 ± 0.5 ‰ | -25.1 ± 0.4 ‰ |
| **roots** | -25.4 ± 0.2 ‰ | -25.0 ± 0.3 ‰ | -25.3 ± 0.2 ‰ | -25.2 ± 0.4 ‰ |

**Supporting equations**

The following equations were used to calculate leaf gas exchange parameters according to (von Caemmerer & Farquhar, 1981):

$$E=\frac{u}{s}\times\frac{\left( w_{o}-w_{e} \right)}{\left( 1-w_{o} \right)} \left( Eqn. S1 \right)$$

where E is the transpiration rate in mmol m^-2^ s^-1^, u is the molar flux in mol s^-1^ of air through the cuvette, s is the leaf area in m^2^ and w_o_ and w_e_ are the mass fractions of water vapour exiting and entering the cuvette [mol mol^-1^], respectively.

$$A_{\mathrm{net}}/ R_{n}=\frac{u}{s}\times\left( \frac{1-w_{e}}{1-w_{o}} \right)\times\left( c_{e}-c_{o} \right)-E\times c_{e} \left( Eqn. S2 \right)$$

where A_net_ and R_n_ are the diurnal CO_2_ assimilation and the nocturnal CO_2_ respiration rate [µmol m^-2^ s^-1^], respectively, u is the molar flux in mol s^-1^ of air through the cuvette, s is the leaf area in m^2^ and c_o_ and c_e_ are the mass fractions of CO_2_ exiting and entering the cuvette [mol mol^-1^], respectively.

$$g_{s}=\frac{E\times\left( 1-\frac{w_{i}+w_{a}}{2} \right)}{\left( w_{i}-w_{a} \right)} \left( Eqn. S3 \right)$$

where g_s_ is stomatal conductance for water vapour in mmol m^-2^ s^-1^ and w_i_ and w_a_ are the mass fractions of water vapour inside and outside the leaf [mol mol^-1^], respectively.

$$\mathrm{WUE}=\frac{A_{\mathrm{net}}}{E} \left( Eqn.S4 \right)$$

where WUE is water use efficiency in µmol C mmol^-1^ H_2_O^-1^ and E and A_net_ are derived from formulas S1 and S2.

Continuously measured BVOC fluxes using the PTR-ToF-MS were calculated using Equation S5, while campaign-wise measured BVOC fluxes using GC-MS were calculated using Equation S6:

$$BVOC = \frac{u}{s} \times(c_{o} - c_{e}) (Eqn. S5)$$

where BVOC is the BVOC flux in nmol m^-2^ s^-1^, u is the molar flux in mol s^-1^ of air through the cuvette, s is the leaf area in m^2^ and c_o_ and c_e_ are the mass fractions of BVOC emissions exiting and entering the cuvette [mol mol^-1^], respectively.

$$BVOC = \frac{\frac{(v_{o}- v_{e})\times\frac{u_{e}}{u_{o}}}{s}}{M} (Eqn. S6)$$

where BVOC flux is the BVOC flux in nmol m^-2^ h^-1^, v_o_ and v_e_ are the BVOC concentrations exiting and entering the cuvette [ng], u_o_ and u_e_ are the air fluxes exiting and entering the cuvette [ml min^-1^], s is the leaf area [m^2^] inside the cuvette and M is the molar Mass [g mol^-1^] of the specific compound.

Isotopic ratios (ẟ^13^C) are expressed as the relative deviation from the international Vienna Pee Dee Belemnite Standard (VPDB, Equation S7):

$$\delta^{13}C = (\frac{R_{sample}}{R_{standard}}) -1 \times1000 (Eqn. S7)$$

where R_sample_ is the ratio of ^13^C to ^12^C in the sample and R_standard_ is the VPDB standard.

**Supporting figures**

Figure S 1: Water use efficiency (WUE) of saplings of F. sylvatica (A) and P. menziesii (B) under control (25°C, n=4) and heat stress conditions (35°C, n=8). Points show the-hourly mean of all measured individuals, while lines are fitted with a loess smoothing spline with a span of 0.08. Grey shaded area in the background represents the heat stress period, while the black line shows, when the ^13^CO_2_ pulse label was applied.

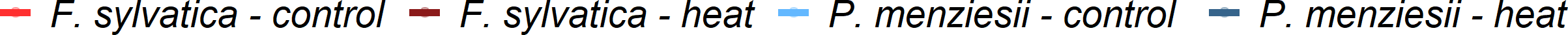

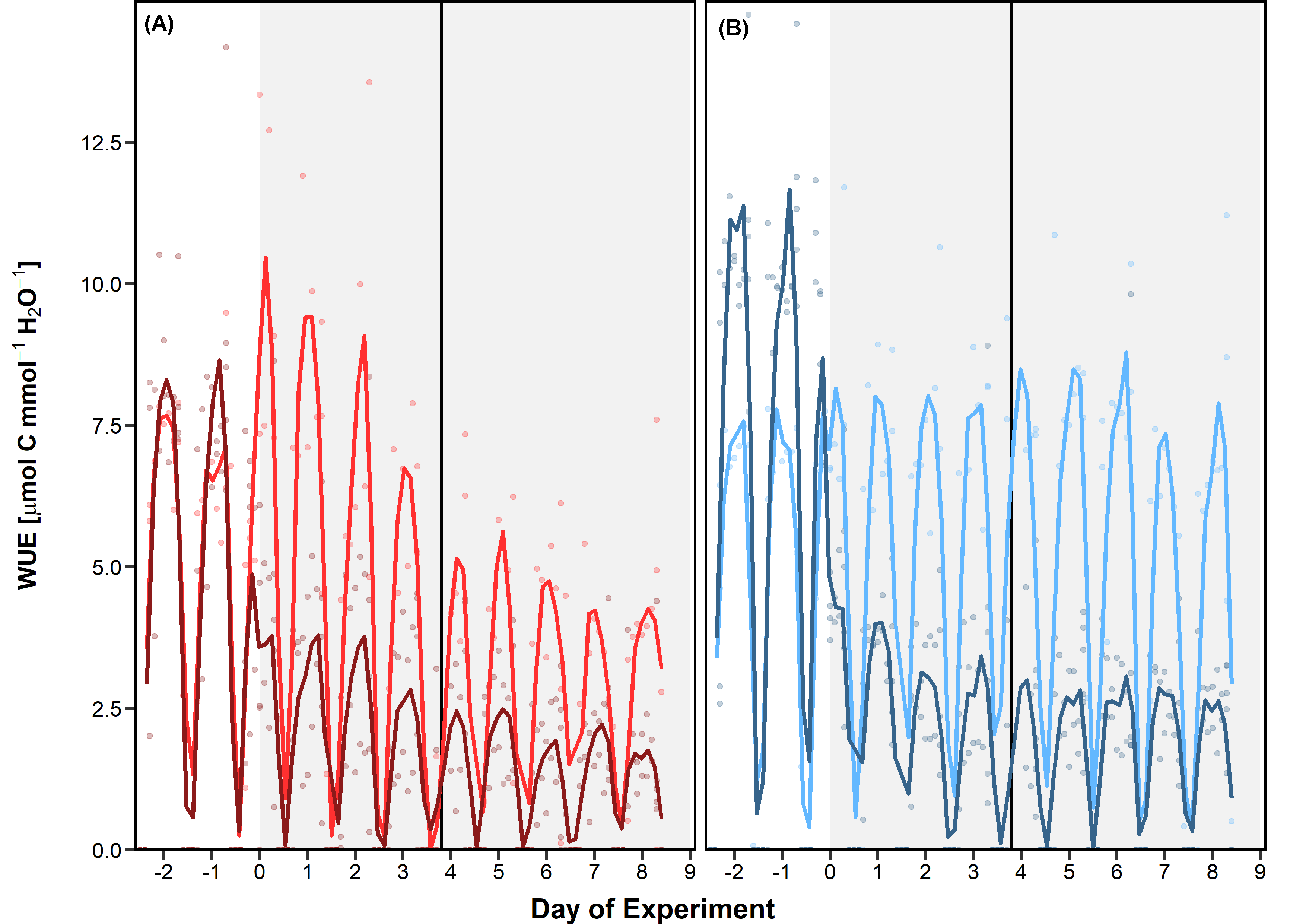

Figure S 2: Water use efficiency of
